# Evidence for convergent evolution of host parasitic manipulation in response to environmental conditions

**DOI:** 10.1101/211144

**Authors:** Raquel G. Loreto, João P.M. Araújo, Ryan M. Kepler, Kimberly R. Fleming, Corrie S. Moreau, David P. Hughes

## Abstract

Environmental conditions exert strong selection on animal behavior. We tested the hypothesis that the altered behavior of hosts due to parasitic manipulation is also subject to selection imposed by changes in environmental conditions over time. Our model system is ants manipulated by parasitic fungi to bite onto vegetation. We analyzed the correlation between forest type (tropical vs. temperate) and biting substrate (leaf vs. twigs), the time required for the fungi to reach reproductive maturity, and the phylogenetic relationship among specimens from tropical and temperate forests in different parts of the globe. We show that the fungal development in temperate forest is longer than the period of time leaves are present and the ants are manipulated to bite twigs. When biting twigs, 90% of the we examined dead ants had their legs wrapped around twigs, which appears to provide better attachment to the plant. Ancestral state character reconstruction suggests that the leaf biting is the ancestral trait and that twig biting is a convergent trait in temperate regions of the globe. These three lines of evidence suggest that changes in environmental conditions have shaped the manipulative behavior of the host by its parasite.

## Introduction

Convergent phenotypic adaptations in response to similar environmental conditions are important evidence of evolution by natural selection. Animal behavior is often a labile phenotypic trait that allows the animal to respond to spatial and temporal environmental heterogeneity. As the environment changes, individuals that can modulate their behavior may avoid, acclimate and tolerate adverse conditions. Over evolutionary time this can result in convergent adaptive behavior observed in unrelated organisms facing similar abiotic pressures. For example, the long-distance migration of birds, insects, whales and turtle are triggered by the changes in temperature and photoperiod (1).

In some cases, the behaviors we observe in nature are an adaptation on the part of parasites that have evolved to infect animals and manipulate their behavior as a transmission strategy (2, 3). In these cases, the behavior of the host (*i.e.* its phenotype) is an extension of the genotype of the parasite; a phenomenon known as the extended phenotype (4). The number of examples of parasites that adaptively manipulate the behavior of their hosts has recently escalated, a reflection of the expansion of the field (5). Many aspects have been considered in the study of parasitic manipulation, such as the mechanisms of behavioral manipulation (6–10), the epidemiological significance of behavioral manipulation (11) and the ecological importance of manipulated hosts in the environment (12–14). What has not been examined is whether parasite manipulation of animal behavior responds to changes in the environmental conditions which the host experiences. Since environmental changes are known to result in adaptive shifts of phenotypes, such as animal behavior (15–18), it is reasonable to suppose that the same environment may act as a selective force on the extended phenotypes of parasites inside those animals.

One system where we might expect the environment to play a significant role in behavioral manipulation is the ‘zombie ant’. In this system, many species of parasitic fungi in the complex *Ophiocordyceps unilateralis sensu lato* (*s.l.*) manipulate ants from the tribe Camponotini to climb and bite onto aerial vegetation, attaching themselves to the plant tissue (19). Uninfected ants never display this stereotypical biting behavior. The biting behavior displayed by the infected ants is the extended phenotype of the fungus and has been experimentally demonstrated to be adaptive for this parasite, which has zero fitness if the host falls on the ground or is moved to the forest canopy or inside the ants’ nest (11, 19). The death of the ant, shortly after the manipulated biting behavior, is the end point of the manipulation and marks the transition for the fungus, from feeding parasitically on living tissue to feeding saprophytically on the dead tissue of its recently killed host (9). Besides providing nutrients, the carcass of the ant will serve as a platform for the fungus to grow a long stalk, externally from its dead host, to release the spores (termed ascospores in this group of fungi) that ultimately will infect new hosts (20, 21). Once it starts growing externally to the dead ant the fungus is exposed to environmental conditions outside the body of its host. Fungal development is known to be strongly affected by environmental conditions, notably changes in humidity and temperature (22, 23). Species in the *O. unilateralis* complex have been recorded at latitudes ranging from 34° north (10) to 20° south (11) which implies a wide range of environmental conditions exist in which behavioral manipulation of the ant and the subsequent *post mortem* development of the fungus occur.

Previous observations have suggested two kinds of behavioral manipulation occurring in distinct forest types. In tropical forests, ants infected by species of fungi in the *O. unilateralis* complex are predominantly manipulated to bite leaves (11, 19, 24). By contrast, in northern temperate systems (USA, Japan), manipulated ants have been described as biting onto twigs (10, 25). The seasonal leaf shed observed in temperate forests represents a major difference in comparison to tropical forests, where the majority of the trees are evergreen with leaves present throughout the year. For a parasite that manipulates the host to bite leaves before using the host cadaver as a platform for transmission, the permanence of a leaf as a platform may impact its fitness. Although there are other parasites that manipulate ant behavior, including other group of fungi (26), as well as cestodes (27), nematodes (28), trematodes (29, 30) and flies (31), none of them have been extensively studied as the zombie ants. This deeper understanding of the biology implies the zombie ant system may be a more suitable model for studying how environmental variation affects the behavioral manipulation of hosts by parasites.

We hypothesized that biting different substrates (leaf *versus* twig) is an adaptation of the parasite extended phenotype to the distinct seasonality and environmental conditions present in the two forest types (*i.e.* tropical *vs.* temperate). To test this hypothesis, we focused on three lines of evidence. First, it was necessary to confirm if the biting substrate (leaf *vs.* twig) consistently varies across the South-North cline from tropical forests to temperate woods (in the northern hemisphere). To this end, we analyzed the geographic distribution of species within the *O. unilateralis* complex at the global scale to determine how this distribution relates to the biting substrate. Secondly, we hypothesized that twig biting may confer an adaptive advantage in temperate forests where the leaves are shed annually, especially if the fungus requires an extended period of time to fully develop. Thus, we evaluated, across 20 months, the development of a species belonging to the *O. unilateralis* complex in a temperate forest located in South Carolina, USA after the fungus manipulated its host. Finally, since temperate forests occur in different locations, we tested the hypothesis that behavioral manipulation of ants by fungi to bite twigs is an adaptation that has convergently evolved in geographically distinct temperate forests. To achieve this, we reconstructed the phylogenetic relationships between different species of fungi within the *O. unilateralis* complex that manipulate the host to bite leaves and those that manipulate their hosts to bite twigs, in both Old and New World temperate and tropical forests. Taken together, we present multiple lines of evidence that suggest that in this group, the parasite extended phenotype has responded to long-term changes of environmental conditions by shifting biting behavior from leaves to twigs. Furthermore, the shift in the behavioral manipulation of the host is a convergently evolved extended phenotype in different areas of the globe.

## Material and Methods

### The global distribution of the zombie ant fungi *O. unilateralis s.l.* and variation in biting substrate

In order to report the distribution of species of fungi *O. unilateralis s.l.* complex at the global level, we collected species records from around the world. We searched in museums and herbarium collections, as well as pictures available on the internet (under the terms “*Ophiocordyceps*”, “*Cordyceps*” and “zombie ants”). Additionally, we added records provided by people who directly contacted the authors of this manuscript with pictures of zombie ants that they found worldwide. Furthermore, we used the senior author laboratory collection, which includes samples collected by the authors of this study and other collaborators. This collection also includes specimens donated by the renowned mycologist Dr. Harry Evans, who has worked on *O. unilateralis s.l.* fungi for more than 40 years (20, 24, 32, 33). All samples could be easily ascribed to the *O. unilateralis* species complex which has a very distinctive macromorphology, where the ascospore producing structure (ascoma) distinctly occupies one side of the stalk (hence the epithet *unilateralis*) or the immature stage emerges as a long stalk from between the head and thorax on the dorsal side of the ant (33). For each record, we collected the following information (when available): country, most precise location available (*e.g.* national park, nearest city), geographic coordinates, ant host, biting substrate, collector, year and source. We classified the substrate as “bark” when the host was biting the base or main trunk of the tree, as well as when it was encountered inside fallen logs (which only occurred in Missouri, USA). The substrate “twig” was designated when host ants bite the wooden material of the vegetation other than the main trunk (*i.e.* twigs). We classified the substrate as “leaf” when the host was biting leaves and its variations, such as spines. The “green twig” classification, which only occurred twice (in Costa Rica and Thailand) was designated when host ants were biting early stages of stems which were photosynthetically active (green indicating the presence of chlorophyll *a*) and lacking cambium. For the specimens we genotyped during the study, we visually inspected the substrate in the field before collecting the samples. For the specimens we did not genotype but rather used the genetic data available on GenBank, we relied on the accuracy of the description of the samples in the original publication, as well as the figures that accompanied those publications.

### Post-mortem parasite development in a temperate forest

We hypothesized that the plant substrate the ants are manipulated to bite (leaf *versus* twig) was related to leaf shed. The rationale for this hypothesis is that leaf shed in temperate biomes would limit the available time for the fungus to reach maturity if the ants were manipulated to bite leaves in this environment. To provide support for this hypothesis it was necessary to study the time required for *O. unilateralis s.l.* to develop reproductive maturity *post mortem* of the ant in a temperate wood setting. This study was conducted in a private temperate forest patch located in Abbeville County, South Carolina, USA (georeference: 34.375215, −82.346937) between December 31, 2009 and August 23, 2011. This woodland, owned by one of us, is dominated by deciduous trees which shed their leaves in the fall (Fig. S1A,B). During this time, searching for cadavers occurred each day for 3 hours/day which was possible because one of us lives on the property. All the cadavers of infected manipulated ants (attached to the vegetation) were tagged, photographed and the biting substrate was recorded (n=287). For the newly killed ants attached to the vegetation (n=29), we recorded the phenology of the fungus of 29. The data collection was done during the entire year, including the summer, to capture leaf biting, if it occurred. One of us spent approximately three hours each day searching twigs and leaves for the presence of manipulated ants. The 29 ants were photographed on a daily basis for the first 60 days, and then every 2 weeks to a month, until August 2011.

### Phylogenetic analyses

To understand the evolution of substrate use and if this preference is a monophyletic or a convergent trait, we selected fungal species from as many different geographic locations as we could, to estimate the phylogenetic relationships between taxa. DNA extractions were done following the protocol as previously described (25). Briefly, the genomic DNA was isolated using chloroform and purified with GeneClean III Kit (MP Biomedicals). Many of the specimens in the senior author laboratory collection, collected in 1970-80’s, were dry and degraded, resulting in low quality DNA templates. These samples were excluded from the phylogenetic analyses.

From the genomic templates, four genes were amplified by PCR. We used two ribosomal genes, nu-LSU (954 bp) and nu-SSU (1,144 bp), and two protein-coding genes, RPB1 (813 bp) and TEF (1,012 bp). The cleaned PCR products were sequenced by Sanger DNA sequencing (Applied Biosystems 3730XL) at Genomics Core Facility service at The Pennsylvania State University. The raw sequence reads were manually edited using Geneious version 8.1.8 (34). Individual gene alignments were generated by MUSCLE (35). For this study, we generated 123 new sequences (31 for SSU, 31 for LSU, 32 for RPB1 and 29 for TEF). The alignment of each gene was inspected manually and concatenated into a single dataset using Geneious version 8.1.1 (34). Ambiguously aligned regions were excluded from phylogenetic analysis and gaps were treated as missing data. The GenBank accession number and Herbarium voucher for all the specimens and genes used in this work are listed in Dataset S2. The aligned length of the concatenated four gene dataset was 3,923 bp. Maximum likelihood (ML) analysis was performed with RAxML version 8.2.4 (36) through the online platform CIPRES (phylo.org) (37). The dataset was divided into eight partitions (one each for SSU and LSU, plus separate partitions for the three codon positions of protein-coding RPB1 and TEF) and the GTRGAMMA model of molecular evolution was applied independently to each partition. Branch support was estimated from 1,000 bootstrap replicates. Bayesian phylogenetic reconstruction was performed with MrBayes v3.2.6 (38), applying the GTR model with gamma distributed rates and invariant sites using the same partition scheme as the ML analysis. The analysis was run with four independent chains for 5 million generations, sampling trees and writing them to file every 500 generations. Runs were examined for convergence with Tracer v1.6.0 (39). The first 25% of trees were discarded as burn-in and posterior probabilities mapped onto a 50% consensus tree. In addition, we performed an ancestral state reconstruction (ASR). This analysis was implemented in Mesquite v3.10 (40). Ancestral character states were estimated across our single most likely topology with each taxa coded according to biting location preference (twig, leaf, or trunk). We implemented the Mk1 likelihood reconstruction method (with default settings), which maximizes the probability the observed states would evolve under a stochastic model of evolution (41, 42).

To test for correlation between character states for biting substrate and geographic location (*i.e.* tropical vs. temperate), we implemented a test of dependence of character evolution as implemented in Mesquite 3.2 (43). This analysis tested the relationship between two discrete characters across a phylogeny taking into account branch lengths, develops estimates of rates of changes for the characters and tests for correlated evolution without relying on ancestral state reconstruction (44). To discriminate whether a four-parameter or eight-parameter model is a better fit to the data, a likelihood ratio test statistic was used. In this analysis, the null hypothesis is that the substrate where the infected ant bite is random, rather than the correlated evolution of such trait in response to environmental conditions.

## Results

### The global distribution of the zombie ant fungi *O. unilateralis s.l.* and variation in biting substrate

To understand if environmental conditions play a role in shaping host behavioral manipulation by parasites, we first aimed to assess the biting substrate and its relationship to the distribution of species in the *O. unilateralis* complex. Based on the data we gathered, we determined that species in the *O. unilateralis* complex have been recorded in 26 countries (Fig. 1, Dataset S1). We found reports of zombie ant fungi in North, Central and South America, Africa, Asia and Oceania (Fig. 1, Dataset S1). The latitudinal gradient of *O. unilateralis s.l* is 74°, ranging from 47° North (Ontario, Canada) to 27° South (Santa Catarina, Brazil). Our dataset was constructed based on different sources and methods of collection (see methods) and for this reason we are not able to infer the relative abundance in different locations of the globe. However, in some cases, records came from more detailed studies (*e.g.* (11, 45)), so we were able to estimate the abundance for those specific areas. In this way, we confirmed previous observations that most of the occurrence records for *O. unilateralis s.l.* are from tropical forests. In the tropics, the majority of the records were of ants manipulated to bite onto leaves. Exact numbers of *O. unilateralis s.l.* killed ants encountered was not recorded but it is in excess of 10,000 samples based on 12 years of field work in the Atlantic rainforests of Brazil (11), Amazonian forest of Brazil (24) and Colombia (46) and lowland forests of Peninsular Thailand (19, 45, 47). Although leaf biting predominates in the tropics we know of two exceptions; one in Costa Rica (online record, Dataset S1) and another in Thailand (48). In both cases the ants are found biting chlorenchymous stems (green stems/twigs that are photosynthetically active and lack cambium) and detailed information about behavior and ecology of these can be found in the SI text.

**Figure 1:**
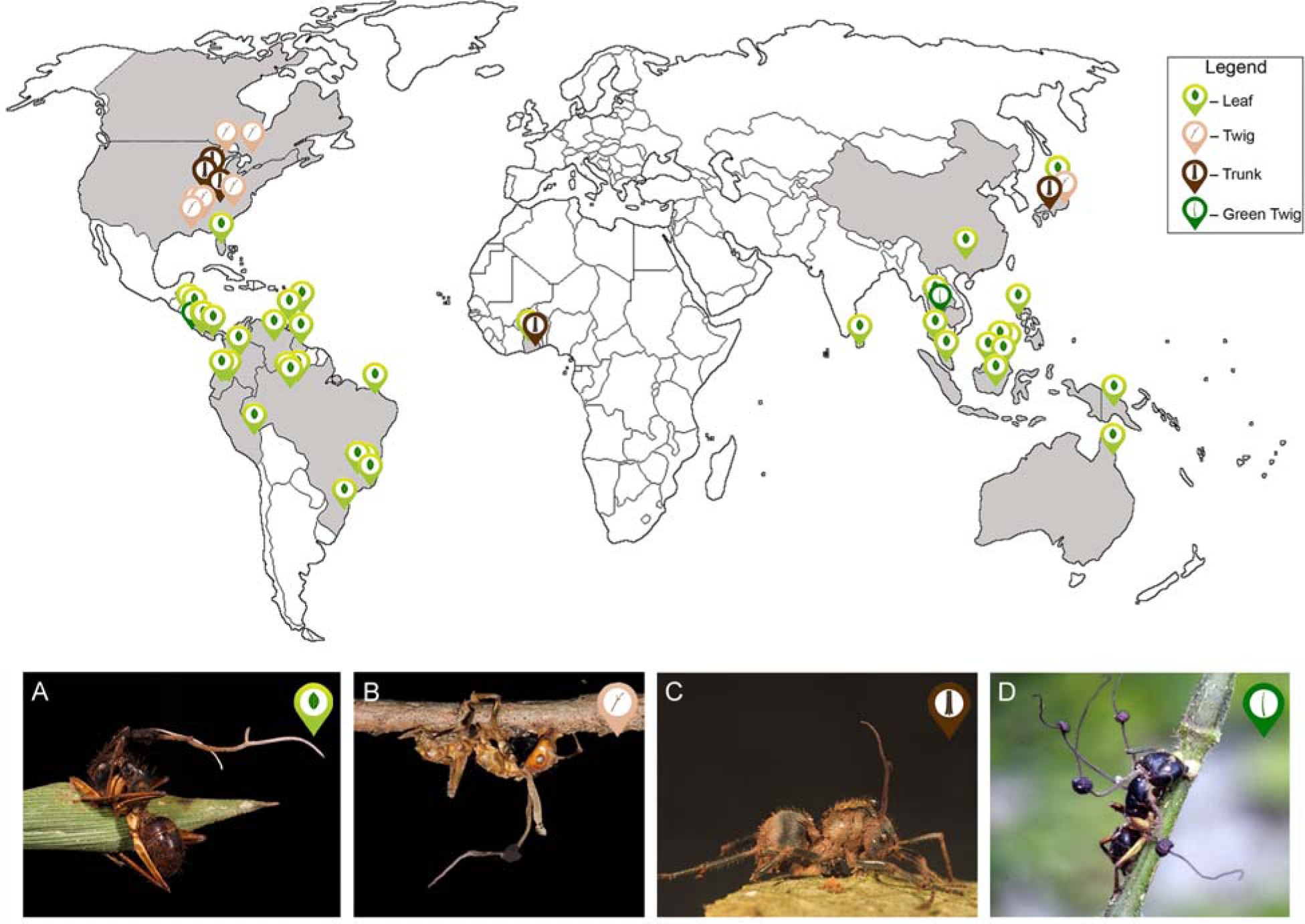
Global distribution and behavioral manipulation by *Ophiocordyceps unilateralis sensu lato* infecting ants. The light green markers represent records of ants manipulated to bite leaves. The light brown represents the records of ants manipulated to bite twigs. The dark brown represents records of ants manipulated to bite tree trunks. The dark green markers represent the record of the one species of ant manipulated to bite green twigs. (A) *Camponotus atriceps* manipulated to bite onto a leaf (Brazilian Amazon). (B) *Camponotus castaneus* manipulated to bite onto twig (South Carolina, USA). (C) *Polyrhachis militaris* manipulated to bite onto bark (Atewa, Ghana). (D) *Camponotus* sp. manipulated to bite green twigs (Nakhon Nayok Thailand), image modified from Kobmoo et al. 2015 (48).

In temperate regions, species in the complex *O. unilateralis s.l.* have been, so far, reported for three countries with predominantly temperate forests: United States, Japan and Canada. In both the United States and Japan, some species of fungi manipulate the ants to bite twigs, while others manipulate their host ants to bite leaves (Dataset S1). In the United States, an undescribed member of this fungal species complex that manipulates the ants to bite onto leaves was reported from an evergreen wetland forest, near the eastern coast of Florida (28° North, Dataset S1). In Japan, we encountered another undescribed species within the *O. unilateralis* complex in temperate forests (30° North), manipulating the ant *Polyrhachis moesta* to bite onto leaves. Interestingly, all the specimens collected for this species were found on evergreen plants (in which there is no leaf fall) in a forest in Kyoto (Dataset S1). In both the USA and Japan, we also encountered ants being manipulated by *O. unilateralis s.l.* to bite onto the bark of trees, although it was not frequent. This bark biting behavior was previously observed for two ant species collected from Ghana, in cocoa plantations (49). In Missouri, USA, we discovered trunk biting where ants were manipulated to bite wood on the inside of logs where carpenter ants had established a colony.

### *Post mortem* parasite development in a temperate forest

We only found cadavers attached to the twigs with the vast majority (286/287) attached to the underside of them (Fig. S1C) and across 20 months we did not find any ants biting leaves. We also found that only two species of ants, *Camponotus castaneus* and *Camponotus americanus* were infected. We conducted extensive searching over the entire year and only discovered newly killed ants between June 20^th^ and October 24^th^ (n=29). Although the newly killed ants were found during the summer and at the beginning of autumn, when both leaves and twigs are available for the manipulated ants to bite onto, all of them were found biting onto twigs. These 29 newly killed ants were labeled with a number and the date they were first found to ensure future identification. For some ants (7/29) we could determine that the cadaver was discovered within the first 24 hours after manipulation and death of the ant, because of the stereotypical appearance of the gaster (terminal portion of the ant’s abdomen) which was noticeably swollen due to abundant fungal tissue inside its body (Fig. 2A, Video S1). The remaining 22 ants were within 2-3 days of death as they were all discovered before the fungus grew from inside to the outside of the ant’s body. As such, all 29 ants were newly killed when first discovered, which provided us the opportunity to record the natural long-term development of the fungus in a temperate forest.

**Figure 2:**
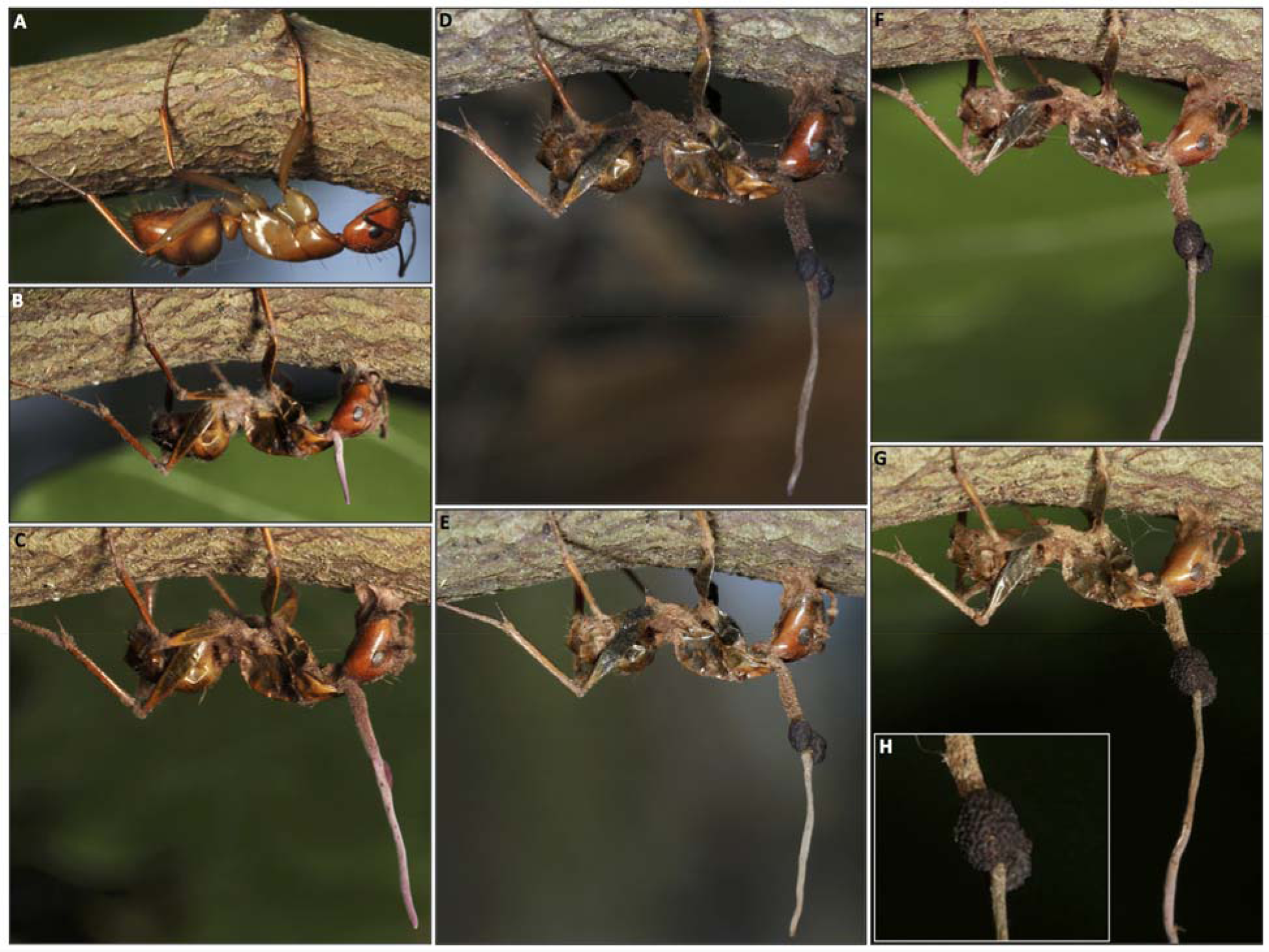
The development of the fungus, *Ophiocordyceps unilateralis sensu lato,* post manipulation and *post mortem* of the ant host, *Camponotus castaneus*, in South Carolina, USA. These photographs were taken of the same ant, under natural conditions, across a year (A) Freshly killed individual (between 0-24h after the ant was killed) on July 25^th^ 2010. (B) August 10^th^ 2010 (C) August 26^th^ 2010 (D) November 14^th^ 2010 (E) March 13^th^ 2011 (F) June 4^th^ 2011 (F) July 16th 2011 (H) Close up of the mature fungal sexual structure.

We discovered that the development of the fungus *post mortem* of the host was delayed until the year following the behavioral manipulation and subsequent death of the ant (Fig. 2). Of the 29 cadavers identified in 2010 (June 20^th^ – October 24^th^) as recently manipulated and killed, 14 fell from the tree soon after biting. The average duration was 25.2 days, ranging from 1-138 days, with the majority (9/14) lasting less than 20 days. Falling may reflect a poor grip on the twigs by the manipulated ant and thus a higher chance the subsequent cadaver would become dislodged. Firmly affixing the manipulated host to the twig by its mandibles is likely difficult because twigs are hard plant tissue that are rounded and larger than the gape of the mandibles. In many samples we encountered, the mandibles did not have a good purchase on the wood. We discovered for ants manipulated by the species of *O. unilateralis* s.l. in South Carolina the behavioral manipulation also involves wrapping the legs of the ant around the twig (Fig. S2 and Movie S2). This is not a behavior observed in healthy ants when they walk or rest on twigs since ants do not walk with their tibia or femurs touching the substrate. Instead, they use the most distal segment of the tarsus: the 3th to 5th tarsomeres (50), each means they essentially walking on their “toes”. Out of the 287 samples we observed during this study, 48 were missing legs, perhaps as a result of a long period in the field. From the 239 of which we could clearly observe details from their legs, 90% (216) had their legs wrapped around the biting substrate. Only 23 (out of 239) ants were attached to the twigs by their mandibles alone. In some cases, as shown in the supporting Movie S2, it is clear that the leg grasping behavior prevented the dead ant from falling from the twig, consequently enabling the parasitic fungus to complete its life cycle. The first pair of legs typically crossed the 2^nd^ and in some cases the 3^rd^ pair of legs, which may provide increased purchase (Fig. S2). Both the legs touching other legs and the legs touching the wood developed dense mats of hyphae at their contact points which stitched the ant to the substrate (Fig. S2). This leg wrapping behavior has never been seen in ants manipulated by *O. unilateralis s.l.* in tropical forests.

We monitored the remaining cadavers (15/29) for fungal growth from the first observation in 2010 and then through the first winter (2010) and then the second winter (2011) until March 2012. The average duration of these cadavers was 572 days, ranging from 510 to 615 days. It is possible to determine if the fungus is sexually mature and thus capable of releasing ascospores because the mature ascoma are recognizable by the erumpent ostioles, which are holes through which ascopsores are released. Based on the morphology of ascoma, the remaining 15 specimens did not reach maturity until the summer of the following year (2011), which can be determined either based on the presence/absence of the ascoma or the erumpent nature of the ostioles on the ascoma (Fig. S3). The minimum time required to reach sexual maturity was 310 days (Oct. 16^th^ 2010 – Aug. 22^nd^ 2011). Note that not all samples reached sexual maturity over the course of this study. In some cases, the stalk broke off (Fig. S4A,B) or hyperparasitic fungi infected *O. unilateralis s.l.* (Fig. S4C) preventing the parasite from reaching sexual maturity. It is notable that during winter the fungus experiences severe weather with snow and ice rain (Fig. S1D,E). Based on our observations, the zombie ant fungi *O. unilateralis s.l.* in North American deciduous forests does not complete its lifecycle before leaf shed in the fall. We suggest this is likely the reason all 287 ants we identified during the summer months (when leaves were present) were manipulated to bite/grasp twigs before being killed by the fungus.

### Biogeography

Both the global dataset we constructed and the phenology study we conducted indicate that biting/grasping twigs is an adaptive trait in temperate biomes due to the fact that twigs are a stable platform for this slow-growing fungus, unlike leaves which are commonly used by infected ants in tropical evergreen biomes. Based on our phylogenetic reconstruction, we find that the species complex *O. unilateralis s.l* forms a monophyletic group (maximum likelihood analysis boot strap value=100). The Randomized Axelerated Maximum Likelihood (RAxML) inference topology is presented in Fig. 3. In addition, we performed a Bayesian analysis with 5 million generations sampled which produced a topology consistent with the ML approach, as well as similar support values for the majority of the nodes (Fig. S5) Within the *O. unilateralis s.l*. clade we recovered two major sub-clades: one formed by Asia-Oceania species and another with mostly American species, with a single exception, *Ophiocordyceps pulvinata* (from Japan). Within the two sub-clades (continental scale), the fungal species did not cluster according to geographic origin of the samples (country scale). Within the Asia-Oceania cluster, where the fungus is found infecting both carpenter and spiny ants, the genera *Camponotus* and *Polyrhachis* respectively, there was also no clustering of fungal species by the ant host although there is host-specificity at the species level. Thus, below the continent level, there is no clear phylogenetic pattern either related to geographic location or the host species.

**Figure 3:**
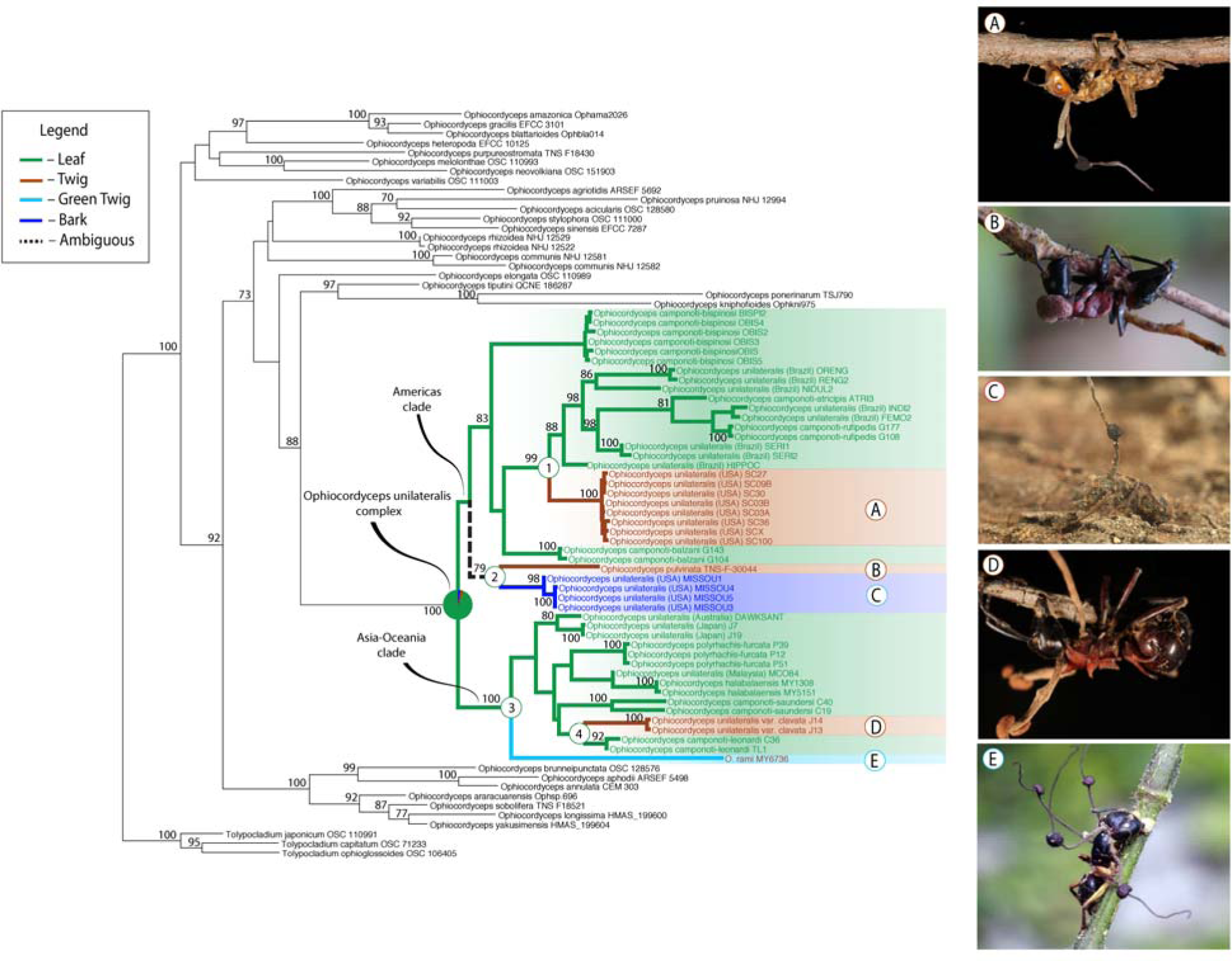
The evolutionary relationships among closely related species of fungi from the *Ophiocordyceps unilateralis* complex that manipulates ants to bite different plant substrates. Phylogenetic relationship between fungi manipulating ants to bite leaves, twigs and bark as inferred with molecular data. (A) *Camponotus castaneus* infected with *O. unilateralis s.l.*, manipulated to bite onto twigs. Samples originated from South Carolina, USA. (B) *Camponotus obiscurips* infected with *O.* (=unilateralis) *pulvinata*, manipulated to bite twigs. Sample originated from Japan; image from Kepler et al. 2011 (25). (C) *Camponotus chromaiodes* infected with *O. unilateralis s.l.*, manipulated to bite onto the interior surface of a tree trunk. Sample originated from Missouri, United States. (D) *Polyrhachis lamellidens* infected by *O. unilateralis s.l.* (var. *clavata*), manipulated to bite twigs. Sample originated from Japan. (E) *Camponotus* sp. infected with *O. rami*, manipulated to bite non-woody twigs. Sample originated from Thailand; image from Kobmoo et al. 2015 (48).

To establish the ancestral condition of biting (leaf or twig), we used Mesquite 3.2 to perform an ancestral character-state reconstruction analysis on our maximum likelihood topology. The result of this analysis is presented in Fig. 3 (colored branches). We find that leaf biting is strongly inferred as the ancestral character state for ants manipulated by fungi within the *O. unilateralis* complex. The shift in biting different substrates occurred at least four times in the evolutionary history of this group of manipulative parasites. In node 1 (Fig. 3), the shift from leaf to twig biting was reported for samples from North America. The ancestral biting substrate for node 2 was ambiguous. This node consists of fungi that manipulate ants to bite onto twigs (from Japan), and tree trunks (from Missouri, USA). Increased sampling is needed to determine if trunk biting evolved from twig biting or directly from leaf biting. It is notable that trunk biting involves manipulated ants biting the inside of logs where ants have established a colony. It may be that the more severe cold in Missouri has shifted manipulation from exposed twigs to more protected logs. Node 3 (Fig. 3) represents the change from leaf biting to green twig, a behavior manipulation found in a fungus species from Thailand. Nodes 1-3 were well supported in the phylogenetic analyses (BS>70). The fourth change on the biting substrate is reported for the fungus species *Ophiocordyceps unilateralis* var. *clavata* (51) (node 4 Fig. 3), which manipulates ants to bite twigs. However, this node is not well supported (BS=48). Tests of correlated characters performed in Mesquite using Pagel’s correlation analysis method (43) strongly supported the correlation between biting substrate (leaf vs. non-leaf substrate) and geographical range of species (tropical vs. temperate) (likelihood difference=5.45429; *P*=0.01). Thus, the substrate where the ants are found attached can be positively correlated with the geographical species range rather than the result of chance.

## Discussion

Convergence of traits in response to similar environment is a central topic in evolutionary biology. We examined the interplay of animal behavior and parasite adaptation by testing the hypothesis that extended phenotypes (*i.e.* manipulation of host behavior by parasites) in phylogenetically distinct parasite species have responded to the environmental conditions experienced by both partners in similar manners. The overlap between the observed biting substrate (leaf/twig) and forests system (tropical/temperate), together with the time required for the fungus to complete its life cycle in each system, as well as the homoplasic nature of the biting substrate trait, suggests that environmental conditions have played an important role in shaping the mode of behavioral manipulation by this group of fungal parasites. Based on the ancestral state reconstruction analysis we suggest that leaf biting, and not twig biting, is the ancestral condition and that twig biting evolved independently in multiple temperate forest biomes due to local environmental conditions and resource availability (*i.e.* ephemeral leaves as platforms versus stable twigs). Additionally, given the apparent difficulty in biting twigs we suggest that in temperate systems twig grasping evolved in addition to the biting behavior, which arose first.

What are the evolutionary pathways that lead to the current distribution of the observed pattern of twig biting/grasping in the temperate forests and leaf biting across the tropical forest belt? In this study, we focused on the phylogenetically distinct species of fungi in the *O. unilateralis* complex. This complex occurs within the genus *Ophiocordyceps* which is one of the most speciose taxa of fungal parasites infecting insects (52, 53). We know that fungi in the order Hypocreales, which *Ophiocordyceps* belongs to, were ancestrally associated with plants and transitioned from plant-based nutrition to animal parasitism about 150 million years ago (54). Tropical Asia is likely to be the center of origin of this entomopathogenic fungi group (55) and so they likely originated in moist, lowland tropical forests. The precise age of the *O. unilateralis* clade is unknown, but based on the chronogram of Sung et al. (54) it is likely the early Eocene, 47-56 million years ago. Our phylogenic analysis and ancestral state reconstruction analysis demonstrate that twig biting/grasping is not restricted to a single clade (Fig. 3). Rather, twig biting/grasping arose independently multiple times in the evolutionary history of this fungal group. Given the early Eocene origin of the *O. unilateralis* clade, this means that the group evolved in an ice-free world with high precipitation, average temperatures of 30°C and minimal pole-pole temperature variations (56). It is likely then that this fungal group arose in evergreen biomes. Meanwhile, the deciduous forest spread in response to seasonal drought at the late Eocene cooling in the sub-tropics, and later became adapted to the seasonal cold in temperate regions (57). We know from fossil evidence that the highly characteristic pattern of leaf biting induced in ants by species in the *O. unilateralis s.l.* complex was present 47 million years ago in what is modern day Germany, which was then an evergreen biome and 10 degrees further south than its current location (58). Thus, based on past climate and forest type distribution, fossil evidence of leaf biting and our ancestral state character reconstruction, there are grounds to suggest that the species in the *O. unilateralis s.l.* clade originally manipulated ants to bite leaves and subsequently experienced independent convergent evolution on twig biting by different species in response to global climate change and the emergence of the deciduous forests in different areas of the globe. The emergence of the additional grip to the substrate, the twig grasping, presumably came later as it may increase the likelihood that the host cadaver, which the fungus requires for reproduction, stays in position over extended periods of time.

Alternatively, the observed patterns of twig biting in temperate regions could be due to adaptive plasticity. The type of biting substrate to which the host bites would then be a plastic trait that responds to the environment inhabited by the host. There are many records of adaptive plasticity as a response to environmental changes (59), including behavioral plasticity (60). If the biting substrate is a plastic trait that responds to the environment inhabited by the host, it would imply either one of the two following possibilities. The first possibility is that all or most of the fungi species in the *unilateralis* group would be able to manipulate its specific ant host to bite both onto twigs and leaves depending on the circumstances. We would then expect that in the tropical forests, at least part of the ants would be manipulated to bite both twig and leaves, or that in temperate forests, some of the ants would be found biting leaves (although it would mean the death of the parasite). However, we only know of one possible record of plasticity on the biting substrate which is the *Polyrhachis* sp. being manipulated to bite both onto leaves and bark of cocoa trees in a Ghanaian cocoa farm (49). The second possibility that biting substrate is a plastic trait is that the species of fungi in the temperate forest that have evolved the capability to induce the twig biting, in addition to leaf biting. Unfortunately, we are not able to test this hypothesis since it would require either transplant of fungal/ant species or common garden experiments (cross-infections). These experiments are not possible for two reasons. The first is that these species of fungi are highly specific to the species of ant they infect and they cannot manipulate different species of ants (10). In fact, a comparative genomic analysis of the *O. unilateralis* species isolated from *C. castaneus* (USA, twig biter) and *O.* (=unilateralis) *camponoti-rufipedis*, a species isolated from *C. rufipes* (Brazil, leaf biter) shows that the genes from *Ophiocordyceps* ant-manipulationg fungi which are not shared with other ascomycetes fungi are mostly species-specific (61). The second reason is that ants they infect do not occur in both temperate and tropical forest simultaneously meaning transplant infections could not be carried out. Although we cannot exclude the possibility that plasticity explains the observed pattern of biting twigs in temperate regions, we suggest that convergently evolved extended phenotypes where different ant species are manipulated by different fungal species in different temperate forests (USA/Japan) is likely the most parsimonious interpretation of our data.

Why should most species of zombie ant fungi manipulate ants to bite almost exclusively leaves in tropical forests since tropical forests also have twigs? In tropical forests, twigs are abundant and are likely more stable than the leaves themselves, as the leaves in the tropical forest last between 1.5 months to 4 years depending on the plant species (62). However, the average duration of the cadavers attached to the leaves in tropical environment is approximately five months and where it has been tested the leaves remain for longer than the development of the fungus (45). This implies that in the tropics leaf permanence is not a constraint for the development of the fungus. Perhaps then the preference exhibited by fungal species manipulating ants to bite leaves in tropical forests is related to benefits other than the long-term permanence of the substrate in the environment. It is possible that the underside of leaves provides a favorable microclimate (19, 63), where the developing fungus is protected from UV damage and rain, and experiences more stable temperature and humidity. Compared to the dead tissue of stem bark, the living, vascularized tissue of leaves may also provide a nutritional supplement for the developing fungus. In line with this suggestion, we know that several other entomopathogenic fungi are able to grow as endophytes, including *Beauveria bassiana* (64, 65) in leaves, and *Metarhizium anisopliae* (66) and *Ophiocordyceps sinensis* (67) in plant roots. Leaf biting may create an opening where fungi gain ready access to plant nutrients. In fact, *O. unilateralis s.l*. fungal tissue has been identified inside the damage caused by biting in the leaf tissue in both modern and extinct leaves (58), however the direct interaction between these fungi and the plant substrate remains to be studied. Perhaps then the choice of leaves over twigs by manipulating fungi was adaptive and the switch to twig biting only emerged under the strong selective regime that deciduous plants present.

Why would deciduous forests be such a strong selective force and why might twig biting be adaptive? To complete its sexual reproduction, fungi within *O. unilateralis* complex grow and mature the ascoma, from where the ascospores will be produced and released to infect new hosts. It is a key point in the life cycle of this parasite, which is dependent on the precise location where the ants are manipulated to die (19). However, the pace of fungal development is generally regulated by temperature (22). In the tropics, the manipulation happens throughout the year (11, 45) and parasite development is completed in a few months after it kills the host (24, 45). In the temperate system, although the manipulation occurs during the summer when the temperature is elevated, our phenology study revealed that winter appears to interrupt development and so it takes at least one year for the fungus to complete the sexual cycle. Previous empirical work showed the placement of the ant cadaver on the forest floor resulted in zero fitness for the parasite (19). If the infected ants were manipulated to bite leaves, the cadaver would fall onto the forest floor before the fungus can reproduce. Although the manipulation to bite twigs allows the fungus to avoid falling onto the ground due to leaf shed, almost 50% of the newly manipulated ants disappeared from the twigs, resulting in zero fitness for the parasite. This could be due to either weak attachment or predation. The same happens in the tropics, where ants suddenly disappear from the leaf substrate (45). Despite the possibility of falling or being predated, the fungus clearly increases its chance to survive and reproduce by avoiding the leaves. Thus, besides providing evidence that the environment has shaped the behavioral manipulation of the ants by the parasitic fungi, it is possible to suggest a possible mechanism by which this happens. We suggest that the slow growth rate, likely due to the lower average annual temperature, combined with the leaf fall that occurs between the manipulation and reproduction, selectively favored the fungi to manipulate their host to bite onto twigs. In contrast to the leaves of deciduous trees, twigs last for many seasons, providing a steady platform for the fungus to develop and release spores over extended periods of time. Additionally, the difficulty in biting twigs is likely a selective force for the manipulated ant to grasp the twigs with its legs, a novel behavior not previously observed in ants infected by this group of fungi.

The data presented here provides multiple lines of evidence that suggest a parasitic fungus inside ant hosts can respond to environmental change and alter the way it manipulates its host behavior over evolutionary time. We hypothesize that as the evergreen moist forests of the Eocene, which receded first to drier and then cooler deciduous woods (56), favored the selection of a switch in the manipulative behavior from biting leaves to biting twigs. Twig and leaf biting appears in both America and Asia-Oceania host ant clades, as well as in *Camponotus* and *Polyrhachis* ant hosts. In Ghana, some ant hosts were found biting both leaf and bark (the same ant species was found biting both substrates) (49). This indicates lability/plasticity across evolutionary time that may facilitate switching from biting one substrate to another. It is interesting that in tropical forests, where the abundance (11, 47) and diversity of species from the complex *O. unilateralis* is high (20, 24, 25, 48), the default manipulation is leaf biting. However, twigs are also in abundance on tropical forests. That is, when ants are being manipulated, the environment (a tropical forest) has both twigs and leaves available onto which manipulated ants could bite. Likewise, in temperate systems, the biting occurs during the summer, when the ants also have both leaves and twigs as possible biting substrates. We actively searched for infected *Camponotus* ants attached to leaves in our temperate system and did not encounter any. Therefore, the preference for twigs is not an artifact of leaf fall where only ants biting twigs remain to be sampled. A tantalizing question is which factor may have led the fungi to switch their biting substrate. There is some indication that the microclimate is important for the fungal development (19, 68). Therefore, small changes in temperature, humidity and/or CO_2_ concentration could have been used as clues; but we can only speculate. Although it remains to be discovered how a microbe inside the body of its host can affect such precise choices in its manipulated host, our data suggest that the infected manipulated ants have a behavior, the extended phenotype, which is encoded by the fungus and results in the optimal selection of the plant tissue (leaf *versus* twig) to bite before being killed by the parasite.

## Acknowledgements

We are thankful to Dr. Takuya Sato and Mr. Shigeo Ootake for the hospitality, guidance and help with the field work in Japan. We thank Dr. Harry Evans for providing us access to the samples he collected. We are thankful to Djoshkun Shengjuler for revising our manuscript and to the anonymous reviewers for the comments that improved our work. RGL was partially supported by CAPES-Brazil (grant BEX6203-10-8). This work was supported in part by NSF grants IOS-1558062 (DPH) and NIH grant R01 GM116927-02 (DPH) and a grant from the National Academies Keck Futures Initiative-Collective Behavior: From Cells to Societies program (DPH and CSM).

## Supplementary Information Text

**Additional information on stem biting in tropical forests:** The fungus *O. rami*, which belongs to the *unilateralis* complex (and was formerly described as *O. unilateralis*), was collected in the tropical forests of Thailand (14° north), and originally described to manipulate its host to bite twigs (48). However, from the figure presented by the authors, the substrate onto which the ant was biting was green, and not woody, as we have observed in temperate areas (i.e. chlorenchymous stems lacking cambium). We could not find any further information on the specimen collected in Costa Rica.

**Dataset S1**: Summary of the information collected for *Ophiocordyceps unilateralis s.l.* records from around the world. We searched in museums and herbarium collections, as well as pictures available on the internet (under the terms “*Ophiocordyceps*”, “*Cordyceps*” and “zombie ants”). Additionally, we added records provided by people who directly contacted the author of this manuscript with pictures of zombie ants that they have found worldwide. Furthermore, we used the laboratory collection of senior author, which includes samples collected by the authors of this study and other collaborators. This collection also includes specimens donated by the renowned mycologist Dr. Harry Evans, who has worked on *O. unilateralis s.l.* fungi for more than 40 years. For each record, we collected the following information (when available): country, most precise location available (e.g. national park, nearest city), geographic coordinates, ant host, biting substrate, collector, year and source.

**Dataset S2:** Taxon, specimen voucher and sequence information for specimens used in this study.

**Video S1:** Fungus development from July 24^th^ 2010 to March 12^th^ 2012.

**Video S2**: Importance of the grasping behavior on fixing the cadaver to the twig.

**Figure S1:**
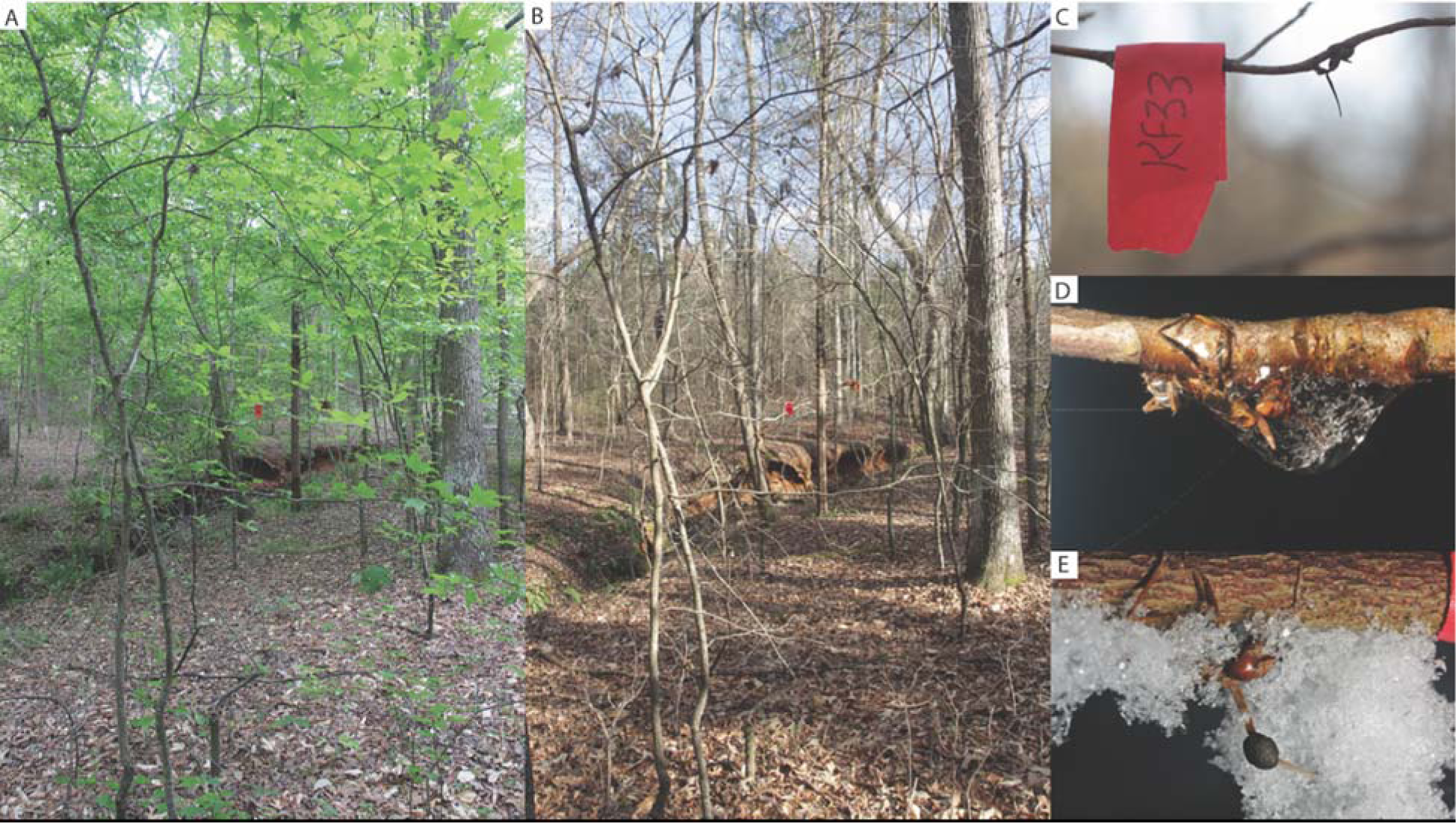
*Ophiocordyceps unilateralis* infected ants in temperate forest of South Carolina, USA. Note that in this natural habitat, the trees display leaves in the summer (A) and shed the leaves in the autumn (B). In this temperate habitat, ants are manipulated to bite the underside of twigs (C). The cadaver of the host and the fungus itself eventually freeze during the winter (D) and/or are covered by snow.

**Figure S2:**
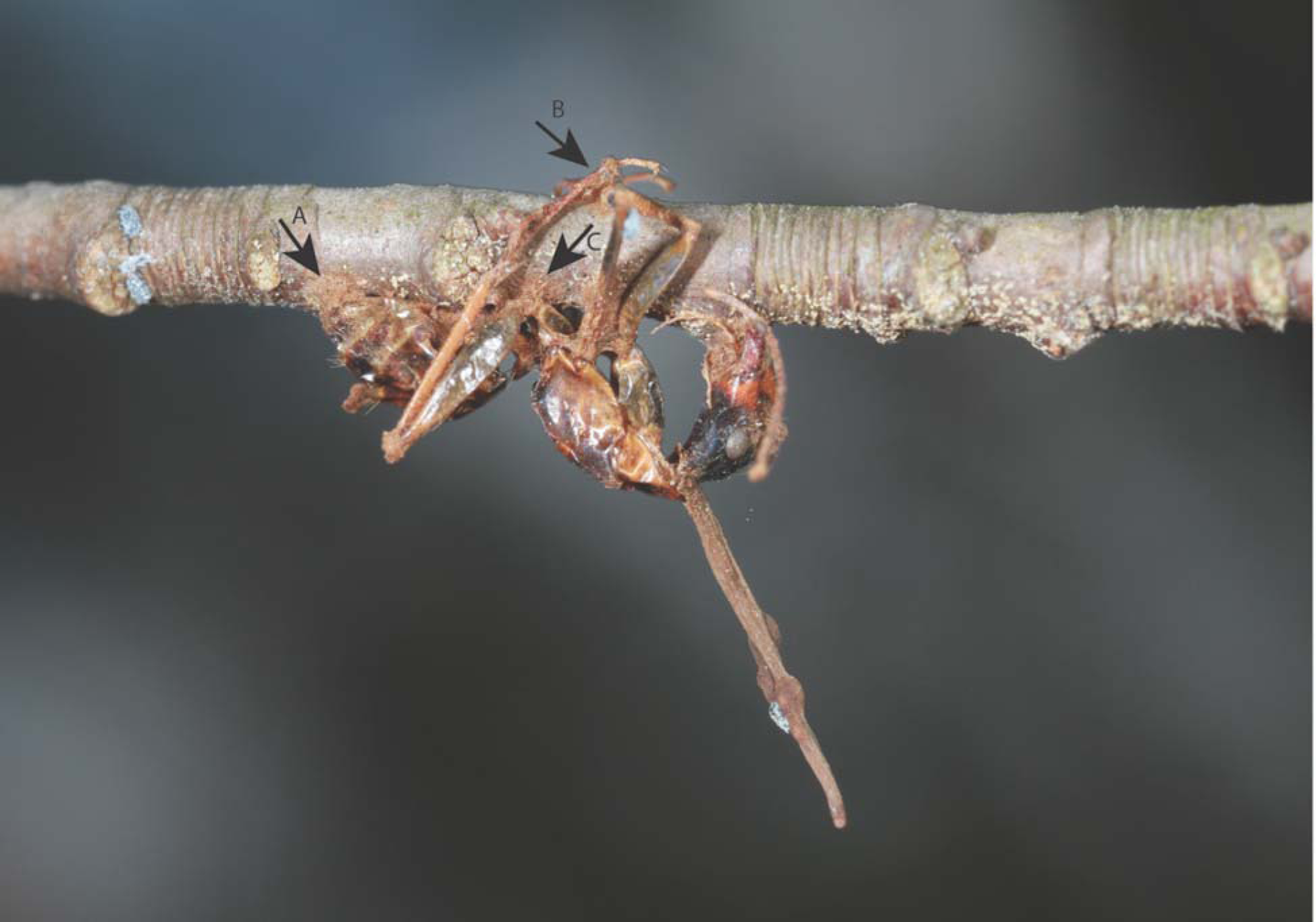
Cadaver of a *Camponotus castaneus* previously manipulated before having been killed by the parasitic fungus *Ophiocordyceps unilateralis*. In this temperate forest, the fungus manipulates the ants to bite and grasp onto the twig. After the death of the host, the fungus develops dense mats of hyphae from the cadaver, at their contact points such as the gaster with the twig (arrow A), the legs with the twig (arrow B) and between the legs that wrap around the twig (arrow C). We suggest that both the grasping and the mats of hyphae are important to keep the cadaver in place, which is required for the long period of fungus development.

**Figure S3:**
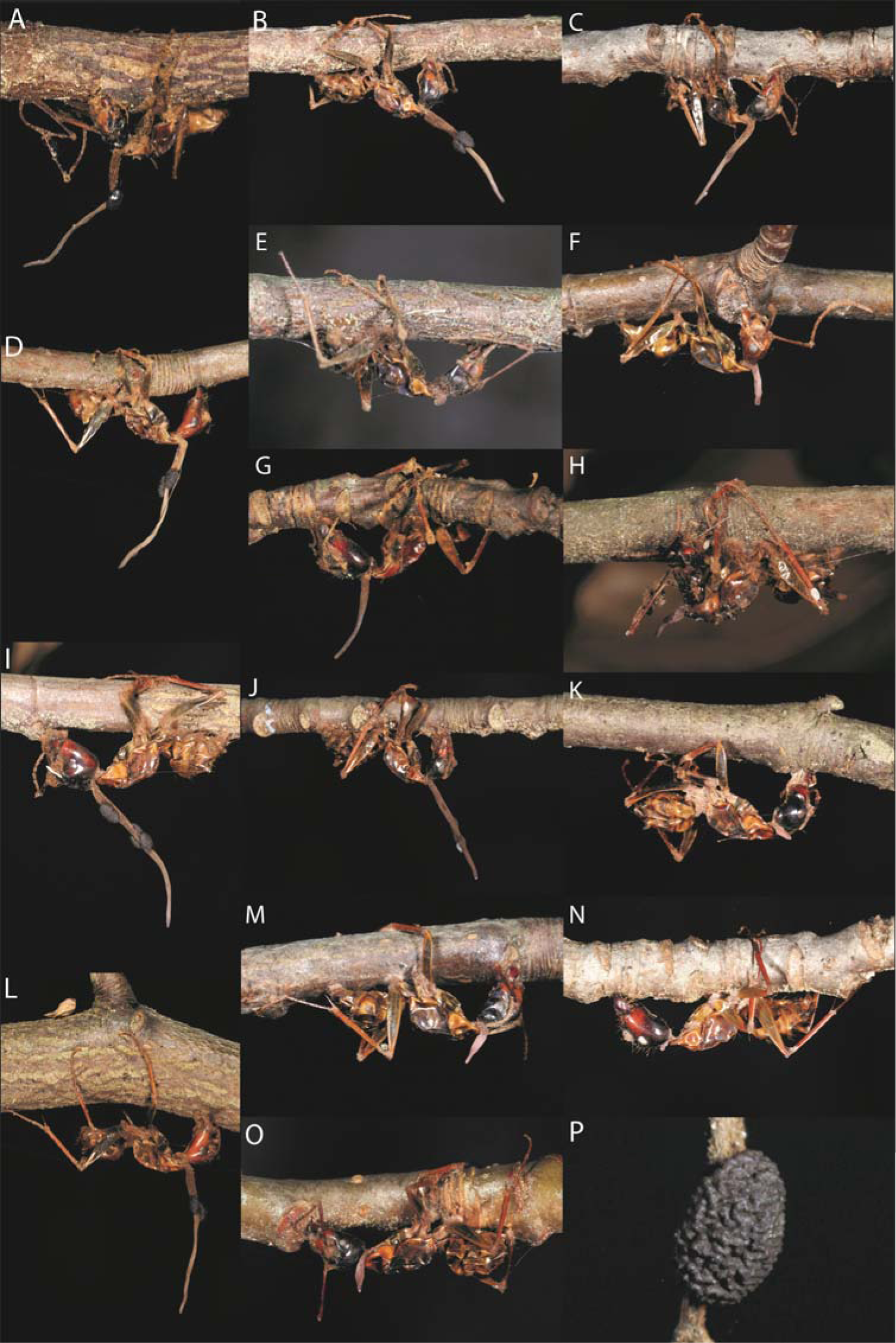
*Ophiocordyceps unilateralis* development. Images A-O shows the fungus development prior its first winter port-mortem of the host for the 15 samples we evaluated in this study. Image P is a close up of a fully developed and mature ascoma. In this study, the morphology of the ascoma was used to classify the ability of the fungus to release ascospores, which is necessary for transmission. Note that none of the samples showed in A-O reached the developmental stage showed in P.

**Figure S4:**
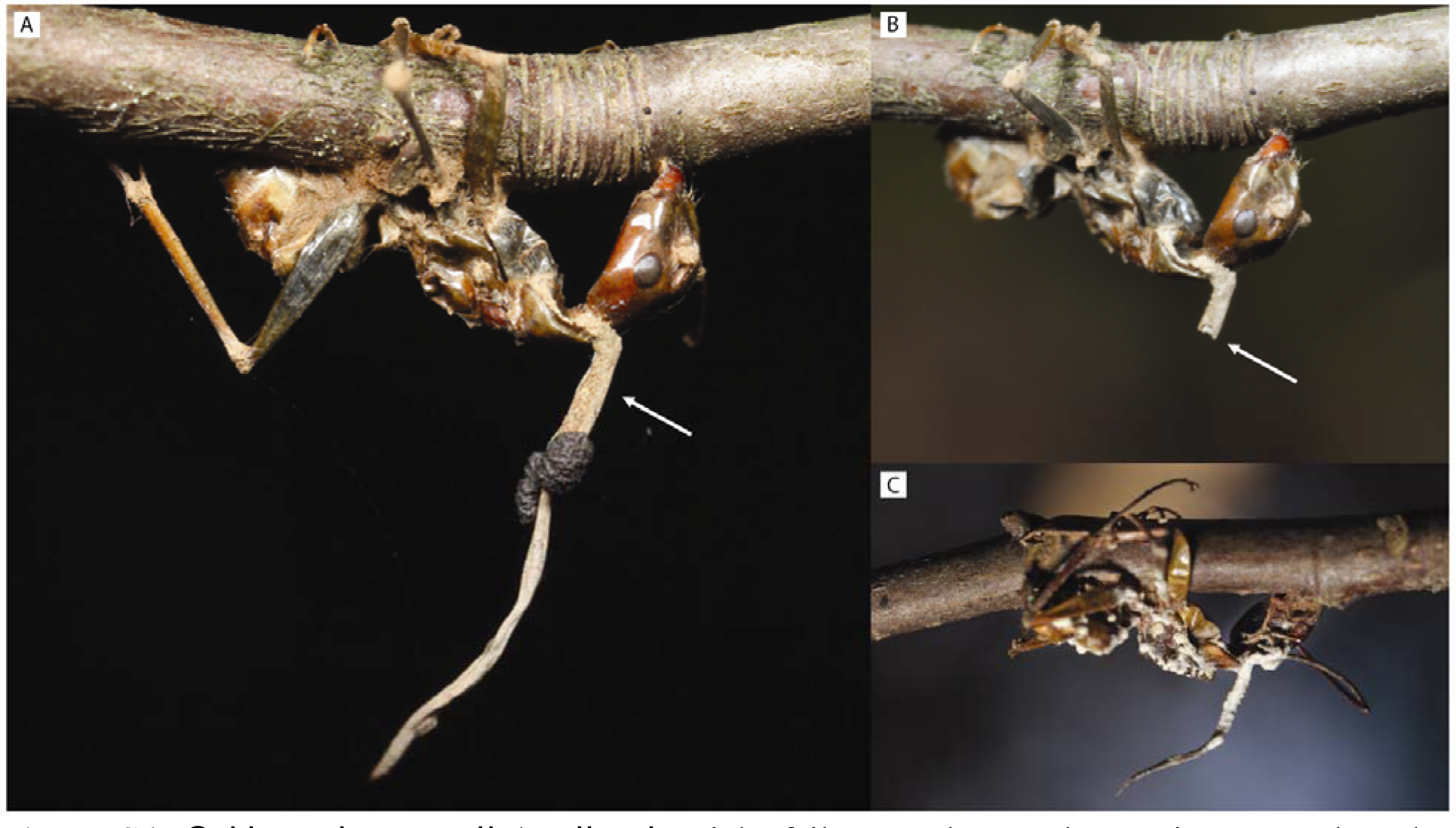
*Ophiocordyceps unilateralis s.l.* might fail to reach sexual maturity even when the cadaver stays attached to the substrate. (A) Sample photographed on January 2011, 5 months after the host’s manipulation and death. (B) Same samples, photographed in March 2011. Although the fungus did develop normally, the stalk broke off, and the fungus did not reach sexual maturity until the end of this study. It is possible that a new stalk and ascoma grew, making it possible for the fungus to sporulate. (C) Specimen of *O. unilateralis s.l.* hyperparasitized by another fungi (micoparasite).

**Figure S5:**
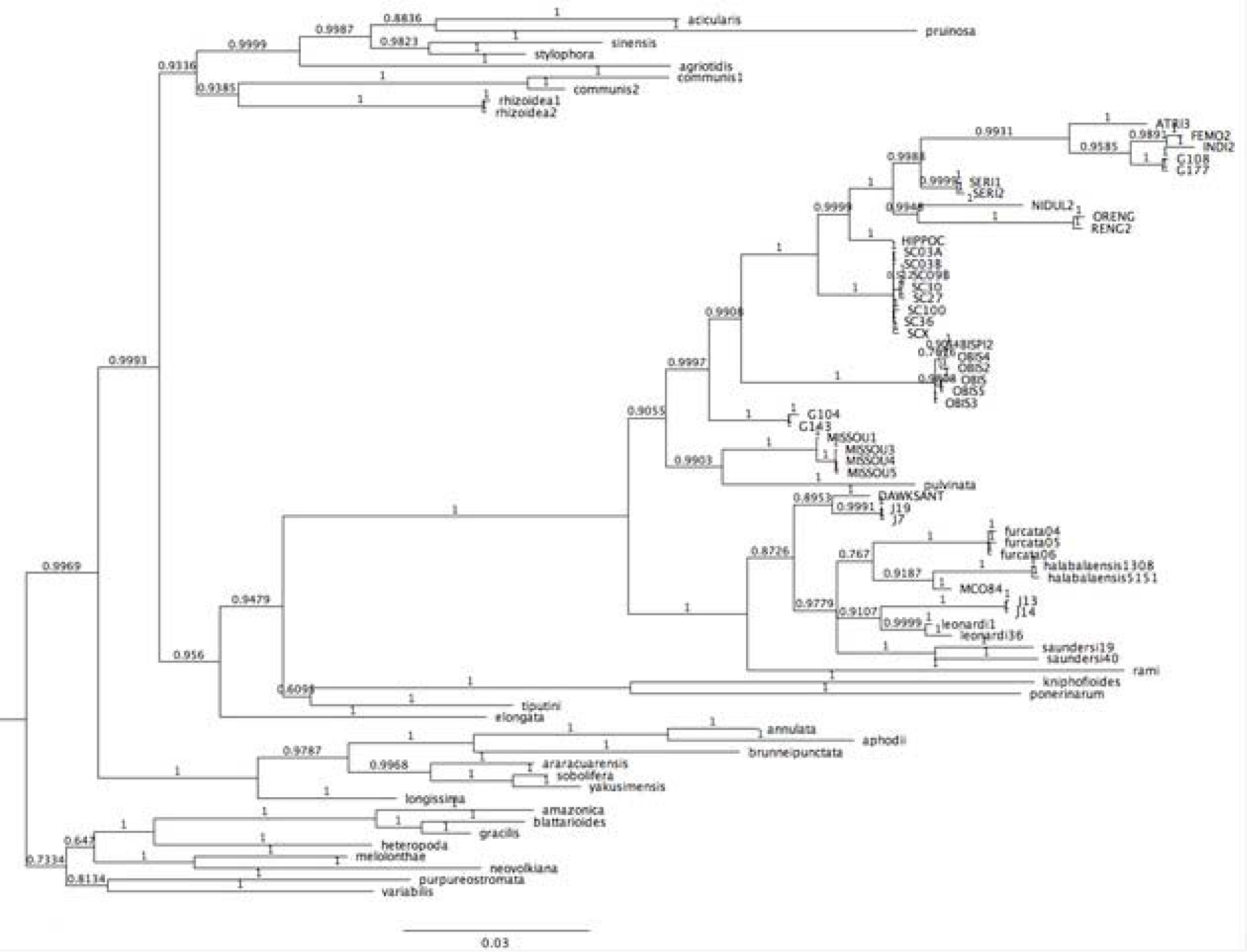
Bayesian tree reconstruction with 5 million generations sampled in MrBayes (using the same model partitions as the ML analysis) produced a topology consistent with the ML approach.

